# Mechanisms of Coupling between Angiotensin Converting Enzyme 2 and Nicotinic Acetylcholine Receptors

**DOI:** 10.1101/2021.08.12.456154

**Authors:** Nadine Kabbani, Kyle D. Brumfield, Patricia Sinclair, Arvind Ramanathan, Menu Leddy, Rita R. Colwell, James L Olds

**Affiliations:** School of Systems Biology, George Mason University; University of Maryland Institute for Advanced Computer Studies, University of Maryland; Maryland Pathogen Research Institute, University of Maryland; Interdiscplinary Program in Neuroscience, George Mason University; Argonne National Laboratory, University of Chicago; Essential Environmental and Engineering Systems; Schar School for Policy and Governance, George Mason University

**Author notes:** **Correspondence:** Nadine Kabbani, 4400 University Drive, Fairfax, VA 22030, USA. NK and KB contributed equally to this work.

**Keywords:** Virus, Toxin, Protein Evolution, Lipid Raft, Angiotensin Converting Enzyme 2, COVID-19, Nicotinic Acetylcholine Receptor, SARS-CoV-2

## Abstract

Severe acute respiratory syndrome coronavirus 2 (SARS-CoV-2), an RNA virus encapsulated by a spike (S) glycoprotein envelope, binds with high affinity to angiotensin converting enzyme 2 (ACE2) during cell entry of a susceptible host. Recent studies suggest nicotinic acetylcholine receptors (nAChRs) play a role in functional ACE2 regulation and nicotine may contribute to the progression of coronavirus disease 2019 (COVID-19). Here, we present evidence for coupling between ACE2 and nAChR through bioinformatic analysis and cell culture experiments. Following molecular and structural protein comparison of over 250 ACE2 vertebrate orthologues, a region of human ACE2 at positions C542-L554 was identified to have sequence similarity to nAChR-binding neurotoxin and rabies virus glycoproteins (RBVG). Furthermore, experiments conducted in PC12 cells indicate a potential for physical interaction between ACE2 and α7 nAChR proteins. Our findings support a model of nAChR involvement in in COVID-19.

## Introduction

In January 2020, an outbreak of SARS-CoV-2 led to the COVID-19 pandemic, resulting in nearly 170M confirmed cases globally by June 2021 [1]. Like other members of the β coronavirus family, this positive-stranded, enveloped RNA virus encodes an S glycoprotein that participates in host cell entry [2]. SARS-CoV-2 S binds with high affinity to ACE2 [3], a transmembrane metalloprotease that cleaves extracellular substrates and is itself subject to proteolytic cleavage [4][5]. While initially discovered as a second form of ACE within the mammalian renin angiotensin system (RAS) [6], orthologs of ACE2 have been reported in numerous species of the subphylum *Vertebrata* [7] and some insects—which do not contain a closed circulatory system [8,9]. Recent work has demonstrated that ferrets, cats, dogs, and select non-human primates are susceptible to SARS-CoV-2 infection [7,10,11]. However, the exact zoonotic tropisms of SARS-CoV-2 in humans remains elusive.

Nicotine exposure, namely through the α7 nicotinic acetylcholine receptor (nAChR), has the potential to promote expression of ACE2 [12,13] and epidemiological surveillance has shown tobacco smokers present a varied response to SARS-CoV-2 infection and outcome [14]. However, the role of nicotine in SARS-CoV-2 infection remains speculative. Differential reports show contrasting lowered and increased number of hospitalizations across subsets of COVID-19 patients with a smoking history [12,15]. The nAChR is a well-characterized protein complex of the cys-loop channel family [16]. Activated by endogenous ligands, such as acetylcholine (ACh), nAChRs are also targeted by many plant and animal toxins [17]. Specifically, α7 nAChR signaling in immune cells can reduce inflammation via tumor necrosis factor α (TNFα) release [18,19]. It is worth noting that nicotine is known to impact the immune system [20], and interactions between nicotine exposure, cholinergic modulation, and ACE2 have been documented within RAS, as well as in neural systems [12,13,21]. ACE2 signaling in neurons and astrocytes can also contribute to oxidative stress and neuroinflammation in the context of Parkinson’s and Alzheimer’s disease [22].

We build on these observations to explore possible sites of interaction between ACE2 and α7 nAChR. Results of molecular bioinformatic analysis reveal a novel sequence domain within human ACE2 position C542-L554 that maintains noticeable sequence homology to several nAChR targeting toxin proteins as well as the rabies virus glycoprotein (RBVG). These findings are corroborated experimentally by evidence of protein coupling between α7 nAChR and ACE2 within cultured cells.

## Methods

### Amino Acid Sequence and Structure Analysis

ACE2 amino acid sequences were compiled from the National Center for Biotechnology Information (NCBI) Reference [23] and the Universal Protein (UniProt) Knowledgebase [24] sequence databases for a total of 256 *Animalia* species of the subphylum *Vertebrata*, using a modified version of a database curated previously [25]. Amino acid sequences of ACE2 orthologues included in this study can be found in **Supplementary File 1**. Amino acid sequences were aligned globally using the MUltiple Sequence Comparison by Log-Expectation (MUSCLE) software package v.3.8.31 with default parameters [26]. A phylogenetic bootstrap consensus tree was constructed from ACE2 alignments using the Molecular Evolutionary Genetics Analysis X (MEGAX) software packages v.10.1.8 [27] employing neighbor-joining method with 1,000 bootstrap replicates, as described previously [25]. Phylogenetic bootstrap consensus trees were visualized using the Interactive Tree of Life web server v.4 [28].

Patterns of ACE2 amino acid conservation across taxa and within accepted phylogenetic relationships were identified with respect to the human ACE2 amino acid sequence (UniProt: Q9BYF1) as reference using Jalview v.2.11.1.0 [29]. Putative common motifs were identified by comparing toxin protein sequences with the following UniProt identifiers (Alpha-cobratoxin, P01391; Alpha-bungarotoxin, P60615; and Long neurotoxin 3P25671) and *Lyssavirus* glycoproteins various isolates (ERA, P03524; CVS-11, O92284; Nishigahara RCEH, Q9IPJ6) to human ACE2 using Clustal Omega v.1.2.4 [30].

Human ACE2 at positions C542-L554, was identified by manually comparing conservation scores with 3D structural models predicted using the SWISS-MODEL web-based service [31] and visualized using PyMol Molecular Graphics System v.2.3.5 (Schrodinger, LLC, New York, NY, USA). To visualize hierarchical clustering of the ACE2 motifs across the species included in this study, a protein BLAST [32] Bit-score ratio (reference/query) was calculated for each protein sequence using human ACE2 as a reference.

### Cell Culture, drug treatment, and microscopy imaging

The pheochromocytoma cell line 12 (PC12) was purchased from the American Type Culture Collection (CRL1721™ ATCC, Gaithersburg, MD, USA) and grown under standard conditions on a collagen (Santa Cruz, Dallas, TX, USA) coated matrix (50 μg/ml). PC12 cells were grown in Roswell Park Memorial Institute (RPMI) 1640 medium (ThermoFisher, Waltham, MA, USA) supplemented with horse serum (10% v/v), fetal bovine serum (5% v/v), and Penicillin-Streptomycin (1% v/v). Prior to experimentation (3 days), cells were differentiated using 200 ng/ml 2.5S mouse nerve growth factor (NGF; Millipore, Temecula, CA, USA) (200 ng/ml), as previously described in [33,34]. To test for protein-protein interactions, cells were treated with the following compounds (reconstituted fresh in serum-free RPMI media): 50 μM nicotine (Sigma), 50 nM α-bungarotoxin, 1 mM choline (ThermoFisher), 1 mM choline (Acros Organics, Geel, Belgium), 15 mM methyl-β-cyclodextrin (MβCD; Sigma Chemical, St. Louis, MO, USA), 5 μM cytochalasin B (CytoB; Enzo Life Sciences, Farmingdale, NY, USA), 50 μM nicotine (Sigma-Aldrich, St. Louis, MO, USA), and 1 μg/ml cholera toxin (*Vibrio cholerae*) (Sigma-Aldrich).

For imaging, cells were fixed using 0.3% glutaraldehyde (80 mM PIPES, 5mM EDTA, 1 mM MgCl2, pH 6.8), probed overnight at 4 °C with polyclonal anti-ACE2 antibody (ARP-53751; Aviva Biosystems, San Diego, CA, USA), and stained with an anti-rabbit secondary fluorescent antibody (Jackson ImmunoResearch, West Grove, PA, USA). The α7 nAChR was labeled using and Alexa Fluor-488 conjugated α-bungarotoxin (fBgtx) (Invitrogen, Carlsbad, CA, USA), as described previously [33]. Cell surface labeling was conducted by applying fBgtx for 15 min at 4 °C prior to fixation. Images were captured using the Zeiss LSM 800 confocal microscope and the Zen software package (Carl Zeiss AG, Oberkochen, Germany). Quantification of fluorescence was obtained using ImageJ (NIH, Bethesda, MD, USA). Colocalization was calculated within regions of interest (ROI) through a dual channel overlay with the JACoP ImageJ plugin v.2.0 [35–37].

### Protein Analysis

For membrane fractions, PC12 cells were solubilized with 0.1% Triton X-100 lysis buffer (Tris HCl, pH 8; 150 mM NaCl; 2 mM EDTA; 10% Glycerol) supplemented with EDTA-free protease inhibitor cocktail (Roche, Penzeberg, Germany), as described in [33]. Protein concentrations were determined using the Bradford Assay (ThermoFisher). Co-immunoprecipitation (co-IP) of the α7 nAChR and its interacting proteins was performed as described in [33], with minor modifications. Briefly, 10 μg of a polyclonal antibody to the α7 nAChR (Alomone Labs # ANC-007, Jerusalem, Israel) was conjugated to Protein G Dynabeads (Invitrogen) by mixing at room temperature (23-25 °C) for 20 min. Beads were then incubated with 1000 μg of crude membrane overnight at 4 °C, as described previously [18]. Protein complexes were eluted from the bead matrix using an LDS Buffer (Invitrogen), following manufacturer’s instructions. Proteins were separated on a 4-12% Bis-Tris PAGE gradient gel (Invitrogen) and transferred to nitrocellulose membranes (Invitrogen). Control experiments for the co-IP experiment were conducted by: 1) loading an equal protein amount onto Protein G beads in the absence of an IP antibody; and 2) using a non-specific (anti-CD11) antibody to test the co-IP under the same condition.

Western blot detection was performed using anti-ACE2 (ARP-5375; Aviva Biosystems) primary antibody and HRP secondary antibody (Jackson ImmunoResearch). SeeBlue protein was used as a molecular weight marker. Proteins were loaded at a concentration of 200 μg/lane. Immunoblots were visualized with the SuperSignal West Pico Chemiluminescent Substrate (ThermoScientific) and imaged using a G-Box Imaging System (Syngene, Frederick, MD).

### Statistical Analysis

Statistical significance was assessed using analysis of variance (ANOVA), followed by a Tukey’s post-hoc test using the R Project for Statistical Computing software package v.4.1.0 [38]. All experimental conditions are reported in comparison to the NGF treated control group, where treatment was with the same vehicle used to deliver the drug. The number of cells was comparable across all conditions: n = 17-20 cells per group. A Pearson’s correlation-based colocalization coefficient [37] was obtained from raw measured signal values expressed as ‘r’ with a range between -1 to +1, i.e., -1 = no colocalization; +1 = full colocalization, [39]. Average band intensity measure using Image J (NIH, Bethesda, MD, USA) was obtained from three separate western blot experiments, and a minimum statistical value of p < 0.01 using a Student’s t-test was considered significant.

## Results

### Human ACE2 residues C542-L554 exhibit sequence homology to nAChR binding proteins

ACE2 is the primary molecular target of the SARS-CoV-2 in mammals and the virus appears to have evolved a strong affinity for ACE2 binding as a likely point of cell attachment [40]. We explored genetic relatedness of the ACE2 protein sequence amongst 256 vertebrate orthologs (**Supplement File 1**). Global ACE2 alignment demonstrated that the protein maintains overall conservation throughout *Vertebrata* evolution (**Supplement Fig. 1**). However, specific amino acid residues responsible for SARS-CoV-2 protein binding, catalytic activity, and intracellular calmodulin (CaM) binding within ACE2 appeared to be differentially conserved across orthologs, suggesting that ACE2 function has potential to vary across organisms (**Fig. 1a; Supplement Fig. 1**).

**Figure 1.**
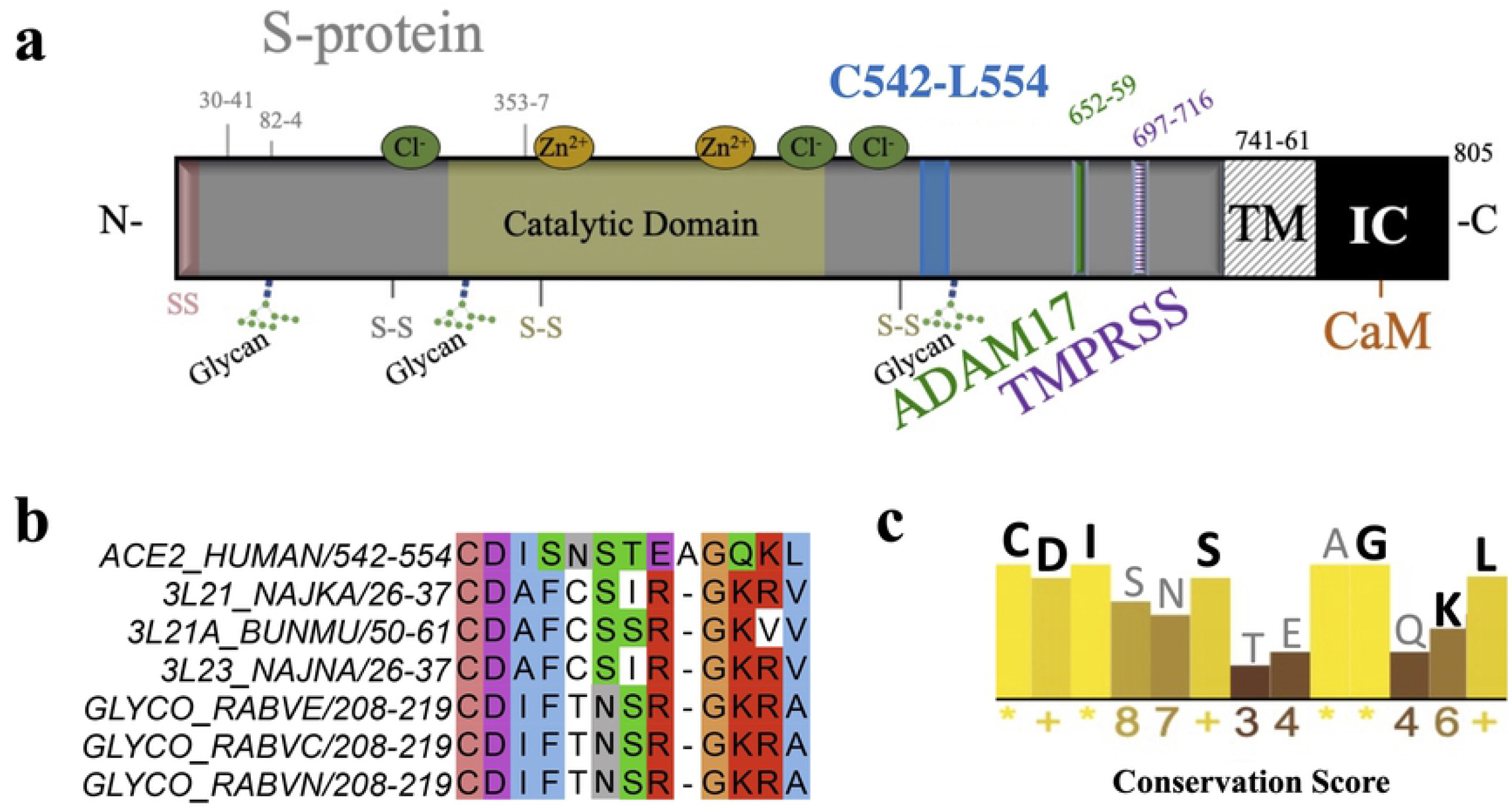
Identification of sequence homology between human ACE2 and nAChR targeting toxin and viral proteins. **a)** schematic of human ACE2 showing residues involved in protein function. **b)** Alignment of human ACE2 (Q9BYF1) residues C542-L554 with neurotoxin and RBVG sequences. **c)** Consensus sequence and histogram depiction of conservation across C542-L554 based on a global alignment of ACE2 vertebrate orthologues. Marks of absolute conserved scores (11) are indicated with a yellow asterisk ‘*’; highly conserved (10) are indicated with a yellow ‘+’; less conserved positions are shown in darker colors.

Earlier research has shown an important role for nAChR signaling in ACE2-mediated regulation of bradykinin and RAS activity within heart, lung, and neural tissue [21], and recent evidence suggests nAChRs participate in inital SARS-CoV-2 infection and subsequent COVID-19 progression [41]. Thus, we sought to test for protein interactions between nAChR and ACE2. Using a multiple sequence alignment algorithm [26], we surveyed for sequence homology between human ACE2 and several nAChR targeting neurotoxin proteins and RBVGs that are known to be homologous to neurotoxins and can bind nAChR in muscle and neural cells [42]. Bioinformatic analysis revealed a specific site of sequence homology between human ACE2, neurotoxin proteins, and the RBVG from multiple rabies virus strains. Specifically, a 13 residue amino acid region within human ACE2 at positions C542-L554 shares sequence homology with neurotoxins and RBVGs (**Fig. 1b**). A comparison of the alignment at the C542-L554 region in ACE2 orthologs shows differential conservation across vertebrates (**Fig. 1c**).

### A structural model of S protein binding predicts that the C542-L554 site in ACE2 is exposed

A recently published cryogenic electron microscopy (cryo-EM) based structural model of human ACE2 bound to SARS-CoV-2 reveals structural regions within ACE2 that participate in interaction with the receptor binding domain (RBD) of the viral S protein (Protein Data Bank: 6MOJ). We used this structure to create a 3D model of human ACE2 using SWISS-MODEL [31]. Such a rendering can explore proximity between various residues and potential sites for protein-protein interaction, based on the 3D protein structure, which cannot be eluciadated from sequence information alone. As shown in **Fig. 2a**, the ACE2 C542-L554 region appear externally on the exposed surface of the human ACE2 protein. Previously identified RBD contact sites of human ACE2 that participate in interactions with SARS-CoV-2 (residues D30-Y41, M82-P84, and K353-R357) [40] were also determined within our structural model (**Fig. 2a**). The ACE2 C542-L554 region is relatively close to the RBD and appears to be accessable by other proteins. Consistent with a published structure showing orientation of human ACE2 when bound to the S protein RBD domain, C542-L554 appears oriented towards the viral protein (**Fig. 2b-c**), and may allosterically contribute to modulating the interaction between the S protein and ACE2.

**Figure 2.**
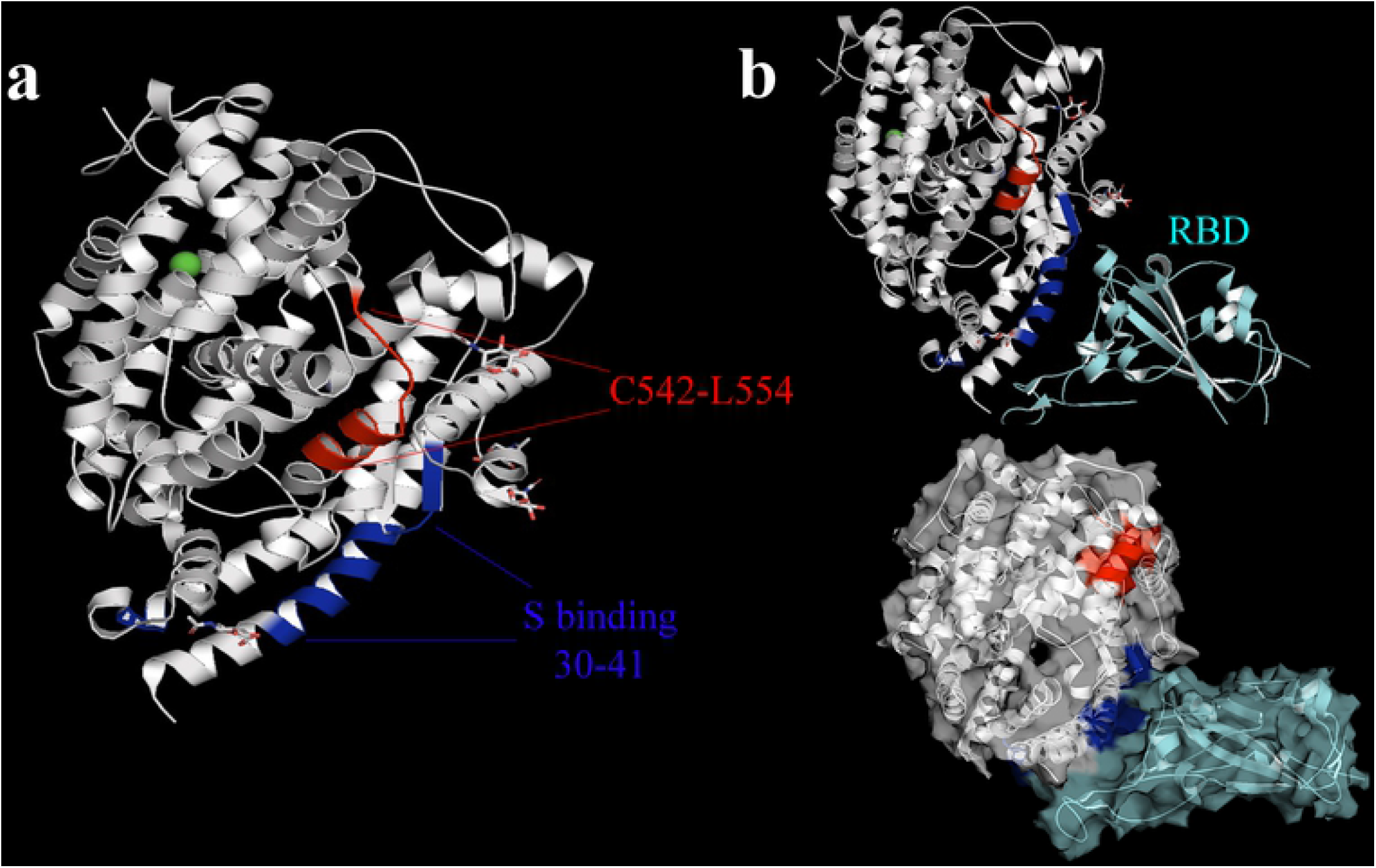
Protein structural model of human ACE2. **a)** Structural rendering of human ACE2 shows the locations of residues C542-L554 (red), and the S protein binding site (blue). **b)** Top: The crystal structure of SARS-CoV-2 spike receptor-binding domain (RBD) (cyan) bound to ACE2. Bottom: surface structures and potential interaction interfaces within the protein complex.

### Conservation and mammalian relatedness of ACE2 across vertebrates

Nicotinic receptors are amongst the oldest channel receptors, with homologs identified in many organisms [16,43]. While less is generally known about the evolution of the ACE2 protein, our findings suggest that it also likely appeared early in vertebrates. We explored evolutionary relatedness among the vertebrate orthologs of ACE2 (**Supplementary File 1**) at C542-L554 using a bootstrapped neighbor-joining method [44]. Phylogenetic analysis of ACE2 proteins from various species suggests that ACE2 sustains common class and taxa properties across vertebrate evolution. This can be seen in **Fig. 3**, which shows that ACE2 shares conservation within all major taxonomic classes, with Aves and Reptilia branching ancestrally to all other classes and forming two major groups. The first includes *Amphibia, Cladistei, Holostei*, and Teleostomi, while Mammalia comprises the second taxonomic group.

**Figure 3.**
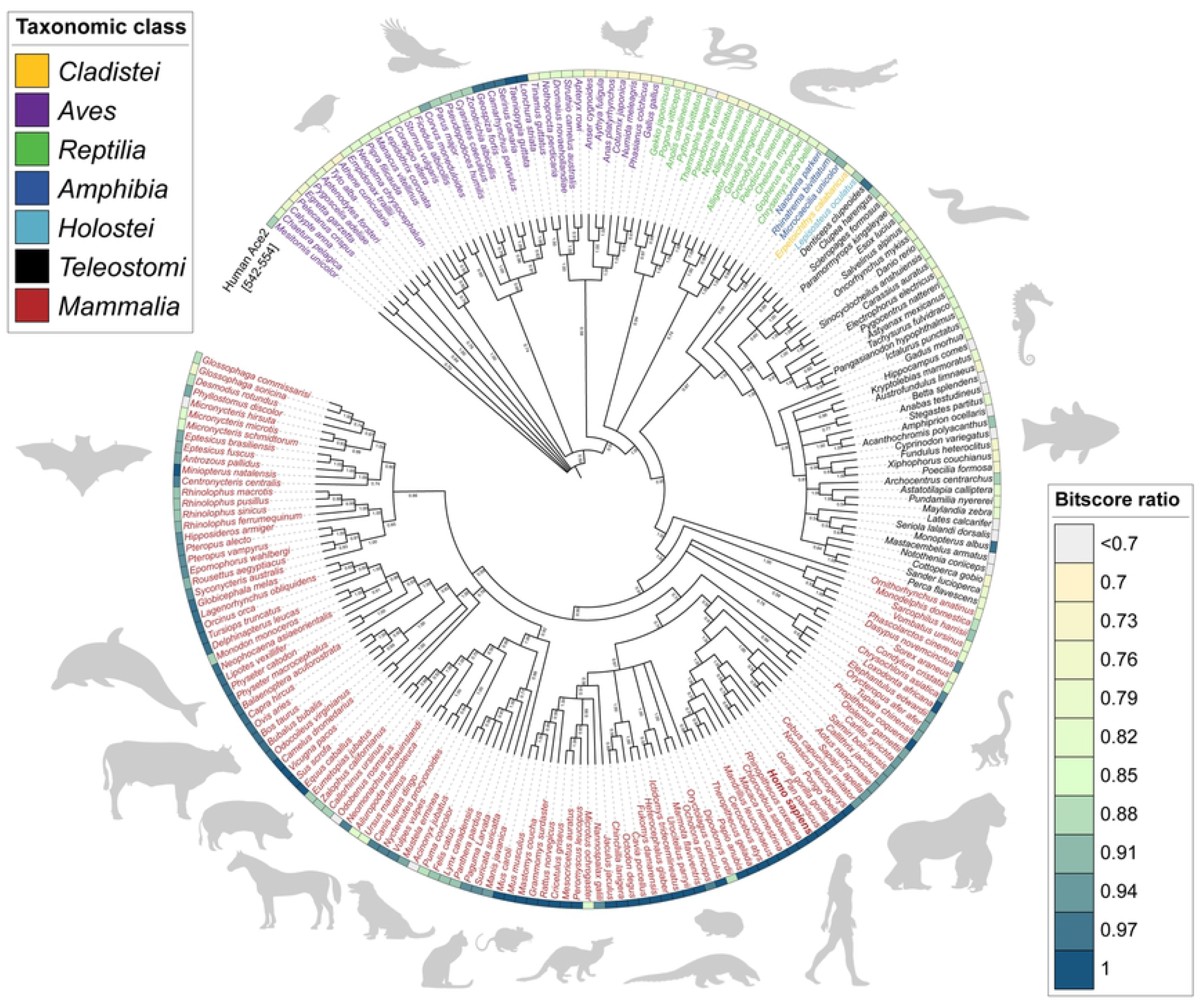
Phylogenetic relatedness of ACE2 orthologues and an analysis of the C542-L554 regions within mammals. An ACE2 phylogenetic tree decorated with a heatmap of amino acid Bit-score ratio similarity between human and other vertebrates. Neighbor-joining bootstrap consensus tree (1,000 bootstrap replicates; >0.5 bootstrap values) shows phylogenetic relationships based on ACE2 amino acid alignment of vertebrate species (yellow, *Cladistei*; purple, *Aves*; green, *Reptilia*; dark blue, *Amphibia*; light blue, *Holostei*; black, *Teleostomi*; red, *Mammalia*). Similarity of human ACE2 (Q9BYF1) C542-L554 against the other vertebrates assessed by BLAST Bit-score ratio (reference/query) is shown as a heatmap encircling the tree.

Conservation was examined across taxa using a protein-protein, bit-score ratio of human ACE2 (C542-L554) as reference. As shown through a heatmap in **Fig. 3**, C542-L554 region appears generally conserved across mammals, with strong similarity between humans and other primates (namely Catarrhines, e.g., Rhesus macaque). In addition, the C542-L554 region is conserved between humans and several other closely related organisms, including *Rodentia* (e.g., house mouse) and *Artiodactyla* (e.g., swine and camel). Interestingly, however, the C542-L554 region showed *less* conservation across bats, with some bats, e.g., *Miniopterus natalensis* (vesper bat), showing strong C542-L554 region homology to human ACE2, while others (e.g., *Micronycteris hirsute* (hairy big-eared bat)) showed low homology.

Generally, birds (*Aves*) did not display homology with humans at the C542-L554 region, with a few exceptions in select species, such as finches. Similarly, most fish (*Teleostomi*) did not show sequence homology to mammals, although, some did show increased homology, e.g., *Monopterus albus* (Asian swamp eel). These results indicate that the human C542-L554 region is conserved across mammals but less conserved across other vertebrates, i.e., *Amphibia, Cladistei, Holostei*, and *Teleostomi*. Species specific variability within several taxa, namely birds and bats, suggests an interesting potential role for this region in ACE2 function and a possible mechanism of SARS-CoV-2 infectivity.

### Protein-protein interaction between ACE2 and α7 nAChRs in PC12 cells

ACE2 is co-expressed with α7 nAChRs in the brain, blood vessels, and kidney and association between the two proteins may contribute to COVID-19 progression [4,5,45,46]. We tested for protein-protein interaction between α7 nAChR and ACE2 within PC12 cells that expresses both proteins endogenously. A co-immunoprecipitation (co-IP) was used to isolate the α7 nAChR-protein from NGF differentiated cells, as previously described [47]. Immunoblot detection showed an association between the α7 nAChR and ACE2 within cells (**Fig. 4a**). Specifically, an anti-ACE2 immunoreactive band was detected in the co-IP experiment at the expected size of full length (rat) ACE2 (∼110kDa). This band was not detected in control lanes where the anti-α7 antibody was omitted or a non-specific antibody was used for co-IP (*see* Materials and Methods). In our membrane fractions, used for both co-IP and western blot detection, several anti-ACE2 immunoreactive bands were detected. Most visible were bands that ran between 100-115 kDa and presumably correspond to the differentially glycosylated ACE2 variants seen in cells [48].

**Figure 4.**
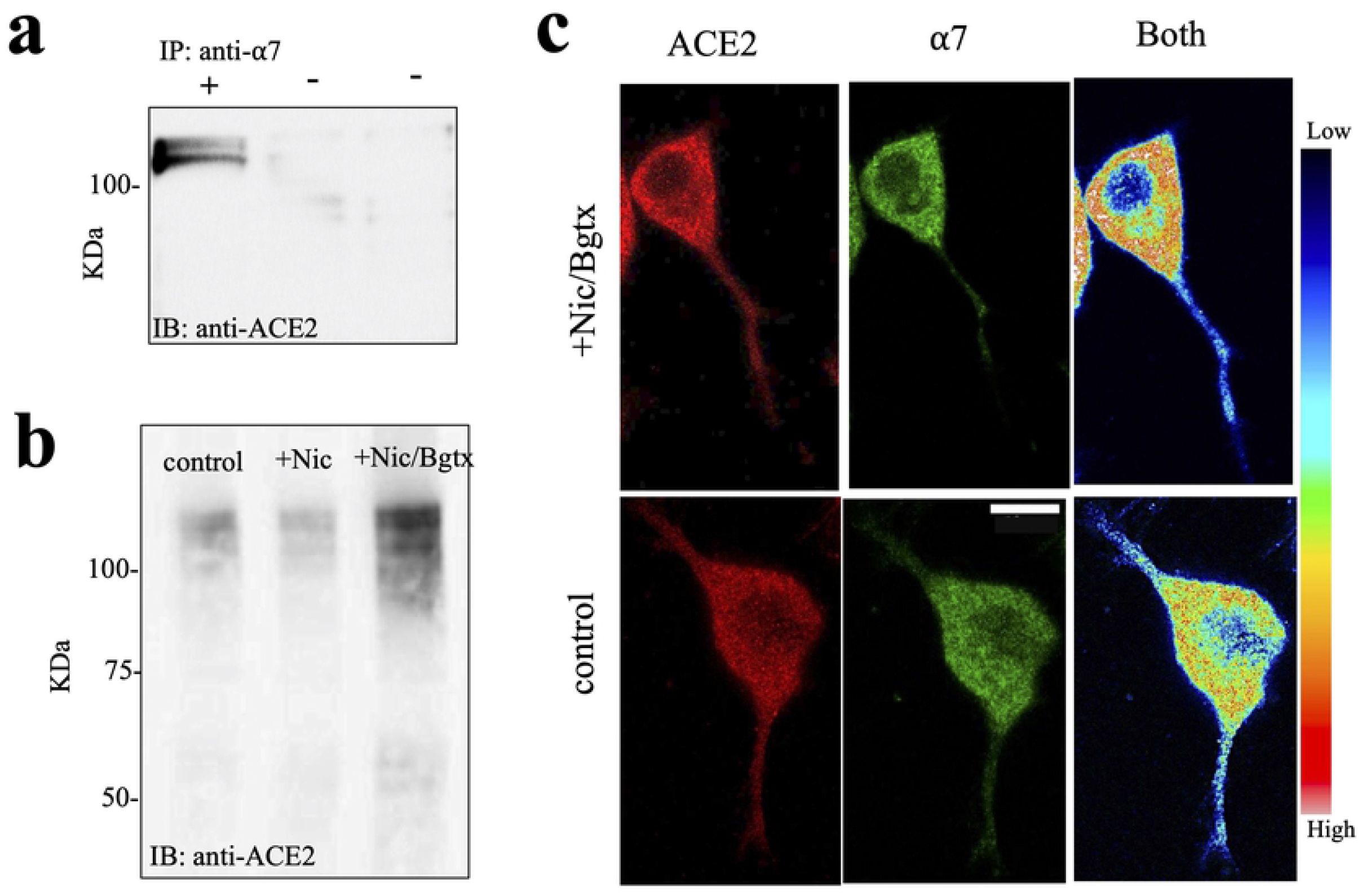
α7 nAChR interacts with ACE2 and regulate its expression. **a)** Co-IP from PC12 cell membrane fractions using an anti-α7 antibody. Western blot detection using an anti-ACE2 antibody shows immunoreactive bands on the gel. **b)** Western blot detection of the ACE2 protein on the gel. **c)** Colocalization of ACE2 and the α7 nAChR (fBgtx) using a heat map to show relative change in the protein colocalization in the cell. Scale bar = 10 μm.

Ligand activation of the α7 nAChR increases intracellular calcium in various types of cells [49,50]. Since ACE2 is calcium regulated through direct intracellular CaM binding, which can participate in ACE2 membrane expression [51] and calcium activation of ADAM17 leading to ectodomain shedding [52], we tested the effect of nicotine agonist activation of nAChRs on ACE2 expression in PC12 cells. Previous studies, suggest that nicotine differentially regulates ACE2 expression, yet this effect appears to vary between cell types [46]. Here, we found that 3-day treatment with 50 μM nicotine decreases expression of ACE2, but this effect was not statistically significant (p > 0.01) (**Fig. 4b**). To test for the specific actions of α7 nAChR activity on ACE2 expression, we co-treated cells with nicotine and the α7-specific antagonist, α-bungarotoxin (Bgtx). As shown in **Fig. 4b**, co-application of Bgtx and nicotine significantly enhanced ACE2 expression on the blot (p < .001).

We examined colocalization between α7 nAChR and ACE2 using immunofluorescence detection of endogenous proteins with fluorescent conjugated Bgtx (fBgtx) and an anti-ACE2 antibodies, respectively. Both signals appear to be broadly expressed in the soma and to a lesser degree in growing neurites (**Fig. 4c)**. At this level of analysis, the protein signals appeared punctate and colocalized intracellularly in a manner consistent with trafficking vesicles. Both proteins also showed some membrane localization consistent with their expected presence at the cell surface. In experiments where nicotine and Bgtx were co-administered, the anti-ACE2 signal appeared more perinuclear (**Fig. 4c**). The evidence points to possible α7 nAChR/ACE2 interaction in the cytoplasm and at the cell surface.

### A role for lipid-rafts in ACE2/α7 nAChR coupling

Cholesterol enriched membrane subdomains, such as lipid rafts, are critical for specificity in protein-protein interactions, and for lipid-associated cell signaling that can contribute to protein turnover [53]. Lipid rafts have also been suggested to be a site for viral cell-surface attachment and for the binding of various toxins [54]. Interestingly, both α7 nAChR and ACE2 appear to localize to rafts [55,56], and thus we tested the involvement of lipid rafts in ACE2/α7 nAChR coupling. In these experiments, cells were treated with choline, a selective agonist of the α7 nAChR [57], and interactions between ACE2 and α7 nAChR were assessed by a co-IP experiment. We compared treatments across three experimental groups: 1) choline alone; 2) choline following the cholesterol disrupting agent MβCD; 3) choline following the inhibitor of actin polymerization cytoB. As shown in **Fig. 5a**, relative to choline alone, ACE2/α7 nAChR interactions appeared stronger when lipid rafts were disrupted. Additionally, ACE2/α7 nAChR interaction appeared dependent on actin assembly. These findings point to cytoskeletal as well as lipid mediated mechanisms that drive ACE2/α7 nAChR binding.

**Figure 5.**
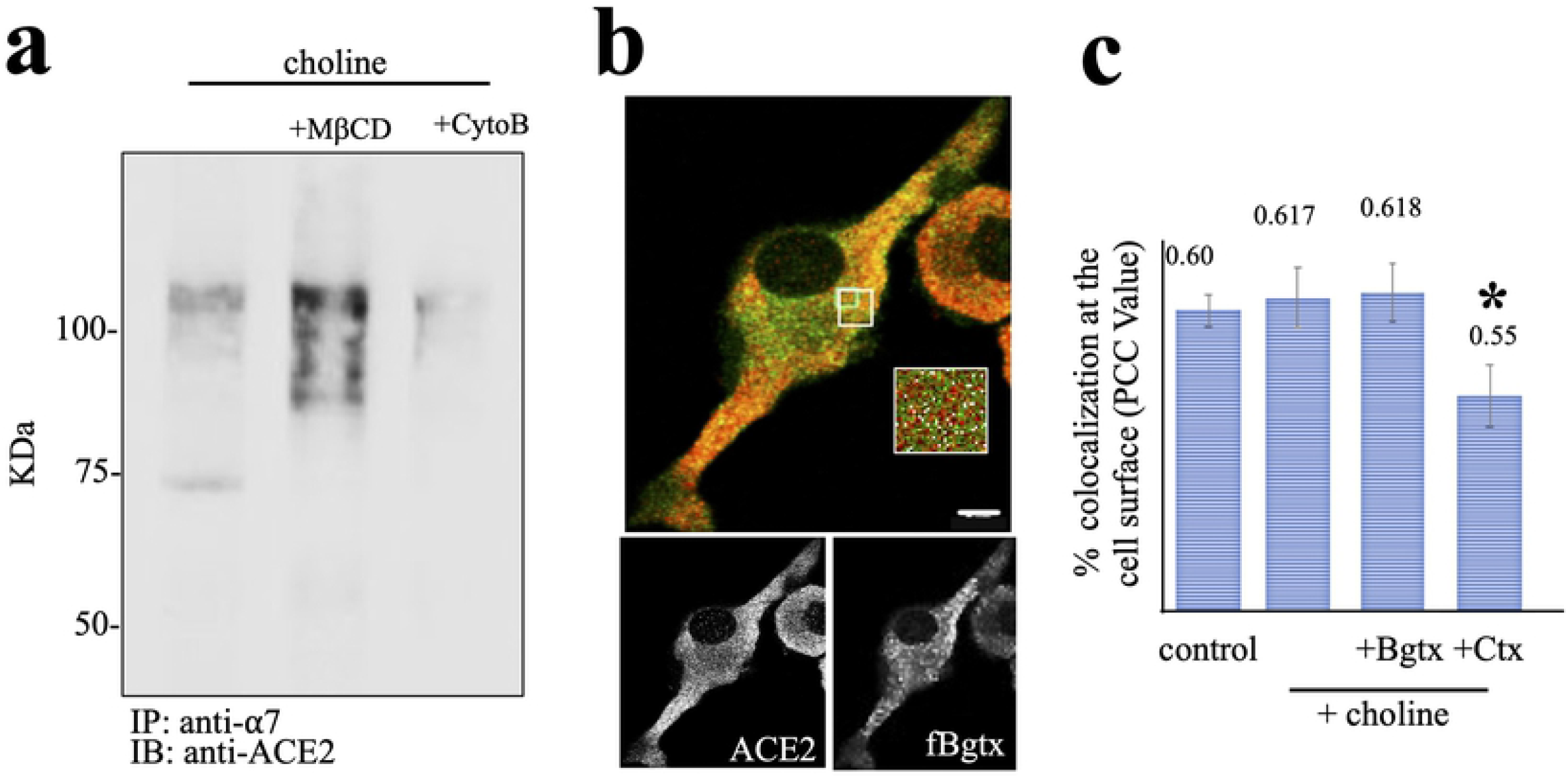
A role for lipid rafts in ACE2 and α7 nAChR association. **a)** Western blot detection of ACE2 protein expression in cells. **b)** Colocalization of ACE2 and cell surface α7 nAChRs (fBgtx). **c)** A Pearson’s correlation coefficient (PCC) of ACE2 and α7 nAChR cellular colocalization with statistical analysis calculated using a one-way ANOVA, followed by Tukey’s HSD post-hoc analysis (F(3,58)=4.194). Scale bar = 5 μm.

We explored colocalization of ACE2 and α7 nAChR at the cell surface using fBgtx to label the receptor prior to membrane permeabilization, and then quantified the extent of anti-ACE2 and fBgtx colocalization. Pearson’s correlation coefficient analysis of colocalization between ACE2 and the α7 nAChR was used to determine the effect of treatment condition across three separate experiments (n=3). As shown in **Fig. 5c,** only cholera toxin (Ctx) treatment was associated with a significant change in ACE2/α7 nAChR colocalization (F(3,38) = 4.194, p = 0.009). In this case, Ctx, which binds to lipid rafts via ganglioside GM1, was associated with a noticeable decrease in co-expression of the two proteins at the cell surface. Based on these findings, ACE2 and α7 nAChR interaction appears to be linked to the lipid membrane in a process that influences protein cell surface expression and potentially internalization.

## Discussion

Evolutionary analysis of sequenced ACE2 orthologs across *Vertebrata* suggests that the C542-L554 region is generally conserved across mammals, yet maintains heterogeneity amongst birds, fish, and reptiles. If this site, indeed, contributes to nAChR binding, our findings support earlier work on involvement of the nicotinic cholinergic system in COVID-19 [41,58,59]. Here, ACE2 targeting may provide a mechanism for viral association with a nAChR multiprotein complex at the plasma membrane, and facilitate viral membrane insertion at regions such as lipid rafts. Indeed, lipid rafts are membrane microdomains that are enriched in cholesterol, glycosphingolipids, and glycosyl-phosphatidylinositol GPI-anchored proteins that have been found to be involved in the replication cycle of many viruses, including coronaviruses [56]. In addition, for many viruses and toxin proteins, cell-mediated attachment and routes of entry appear permissive at rafts [60]. Our experiments show that lipid raft disruption, through MβCD, can impact interaction between nAChR and ACE2. Evidence based Ctx exposure further suggests that nAChR/ACE2 colocalization at the cell surface may also be impacted. Since previous studies have shown that MβCD can interfere with the entry of viruses such as SARS-CoV in the cell [54], it is interesting to consider that interactions between ACE2 and the nAChR represent a convergent mechanism of membrane targeting by viral and toxin proteins [61], with implications for physiological regulation under non-pathological states.

ACE2 is directly regulated by cellular calcium through the presence of a CaM binding “IQ” domain within its intracellular carboxy region (**Fig. 1a**). Elevations in intracellular calcium are reported to promote ACE2 ectodomain shedding through cleavage by ADAM17 [52], and inhibition of ADAM17 appears protective against COVID-19 [62]. In this regard, the α7 nAChR is a calcium conducting channel that can promote local calcium activity through both extracellular and intracellular stores [49]. Determining how calcium-mediated α7 nAChR signaling impacts ACE2 cleavage and shedding is an important next step. Based on the proximity between the ADAM17 and TMPRSS2 cleavage sites and position C542-L554 within ACE2, it is hypothesized that cleavage of ACE2 may facilitate interaction with the nAChR since both proteins are at the cell surface (**Fig. 6b)**. Our experiments indicate several bands on the gel with sizes for full-length and fragmented ACE2. Thus, while the lower weight bands may correspond to cleaved ACE2, at present we cannot conclude this for two reasons: 1. Our results are specific to PC12 cells and expression of proteolytic enzymes, such as ADAM17, have not been determined; 2. We have not yet sampled the extracellular protein fraction for the presence of shed ACE2. Thus, our results suggest a hypothesis whereby ACE2 may interact with nAChRs in either the same or at the suface of another cell if presented in a soluable form (**Fig. 6b**). Here, it is also interesting to consider that soluable ACE2 may provide SARS-CoV2 access to the CNS through nAChR binding in a manner analogous to other viruses [63].

**Figure 6.**
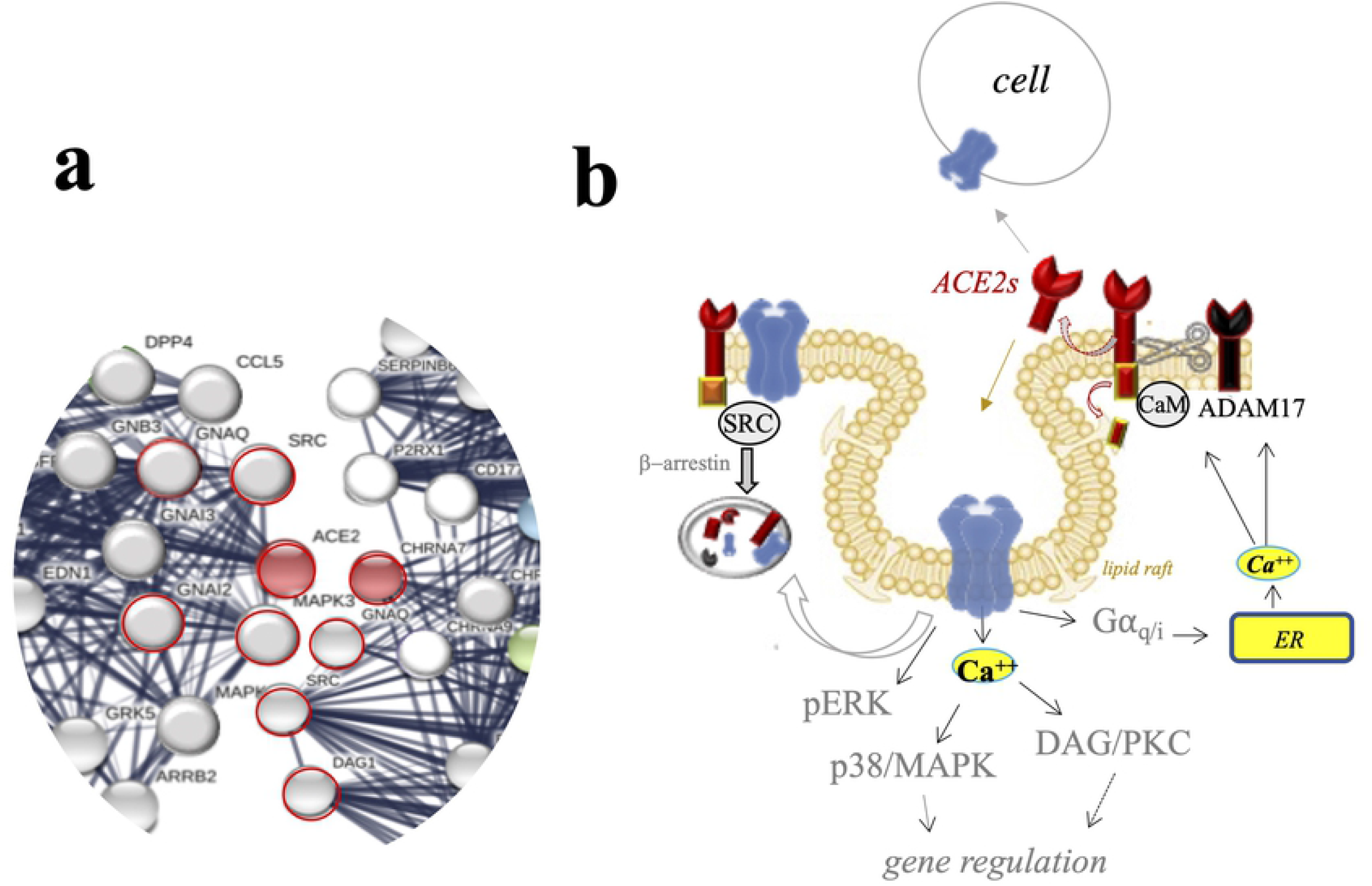
Proposed mechanisms for ACE2 and α7 nAChR coupling in cells. **a)** A STRING database interaction network for α7 nAChR (CHRNA7) and ACE2 shows common pathways (circled in red). **b)** A cell signaling model for ACE2 and α7 nAChR interaction.

Structural findings, based on the 3D model of human ACE2 within PDB [40], suggest that C542-L554 residues face externally within the folded protein and thus may be permissive of protein-protein interaction. These residues appear to also face in the same direction as the region of ACE2 identified for binding the RBD. Hence, while not obstructing the RBD, the location of the C542-L554 site is suggestive of a possible binding site within ACE2 that may modulate interaction with the S protein. In future studies, it will be interesting to determine, both experimentally and computationally, the potential for changes in binding between the nAChR and ACE2 and/or S protein under various states.

Pathogens, namely viruses, share an evolutionalry similarity with biological toxins since both are driven by selectivity for the host. In the example of RBV, nAChR binding enables both acute and long-term neural infectivity in the target organism [64]. The nAChR is also considered to participate in human immunodeficiency virus (HIV) infectivity and cell entry via homology between viral target protein (gp120) and snake neurotoxins [61]. A recent hypothesis for SARS-CoV-2 interaction with nAChR has been proposed [65], and recent simulations suggest nAChRs bind with S protein [41]. However, our data suggests another scenario based on evidence of homology between the ACE2 C542-L554 region with neurotoxin and viral proteins with selectivity at α7 (and potentially α1 and α9) nAChR. Here, the evidence suggests the presnce of *a macromolecular protein complex consisting of ACE2, nAChRs, and the S protein.* This protein complex appears to be functionally linked to the lipid microenvironment of the plasma membrane, where calcium-mediated signaling drives receptor-mediated viral entry and/or vesicular trafficking. Interactome analysis reveals additional pathways of receptor-mediated endocytosis and signaling common to ACE2 and nAChR, including Src kinase and heterotrimeric G proteins (**Fig. 6)**. Our hypothesis mechanism will now need to be tested in future studies that involve computational structural modeling tools and site-directed mutagenesis experiments.

## Abbreviations

nAChRs: Nicotinic Acetylcholine Receptor
ACE2: angiotensin converting enzyme 2
ACh: acetylcholine
Bgtx: α-bungarotoxin
CNS: central nervous system
Co-IP: co-immunoprecipitation
COVID-19: coronavirus disease 2019
Cryo-EM: cryogenic electron microscopy
Ctx: cholera toxin
fBgtx: fluorescent conjugated α-bungarotoxin
CytoB: cytochalasin B
MβCD: methyl-β-cyclodextrin
NGF: nerve growth factor
PC12: pheochromocytoma cell line 12
RBD: receptor binding domain
RBVG: rabies virus glycoprotein
SARS-CoV-2: Severe Acute Respiratory Syndrome Coronavirus 2
S protein: Spike glycoprotein
TNFα: tumor necrosis factor α

## Data accessibility

The authors confirm that the data supporting the findings of this study are available within the article and its supplementary materials.

## Author contributions

NK, KB, AR, JL conceived and designed the project. NK, KB, PS, AR acquired the data. NK, KB, PS, AR, JL analysed and interpreted the data. NK, KB, PS, AR, ML, RC, and JL wrote the paper. All authors contributed to the article and approved the submitted version.

## Acknowledgements

Research reported in this manuscript was supported in part by NIH (R01ES030317A) and NSF Grant (CCF1918749) to RRC.

## Conflicts of interest

The authors have nothing to declare.

## Notes

### Competing Interest Statement

The authors have declared no competing interest.

